# Bacteria-induced colitis in naked mole rats is alleviated by probiotic treatment: a new mammalian model for acute inflammatory disease

**DOI:** 10.1101/2025.06.05.658065

**Authors:** Daniel W. Hart, Aik Seng Ng, Patrycja Gazinska, Robert Goldin, Purva Gopal, Nicolize O’Dell, Shamir Montazid, James E. East, Ian P. Tomlinson, Nadine Koch, Nigel C. Bennett, Shazia Irshad

**Author notes:** Corresponding author: Shazia Irshad. These authors contributed equally to this work (Daniel W. Hart, Aik Seng Ng).

## Abstract

Enteropathogenic bacteria are a major cause of morbidity and mortality globally. Mouse models have been indispensable in advancing our understanding of infectious diseases caused by intestinal pathogens and in identifying physical, chemical and immunological barriers that limit these infections. However, there are significant differences between laboratory mice and human intestinal microbiota and immunobiology that underscore the need to develop other models that recapitulate the disease pathology and mucosal immune responses of human enteric diseases. Here we report how the pathogenic expansion of *Citrobacter braakii* in naked mole rats (NMRs) leads to colonic inflammation and epithelial injury that mimics pathological features of human hemorrhagic colitis. We observe mucosal erosions, ulcerations, depletion of goblet cells, extension of proliferative compartments to the surface of the glands, and active inflammation in the colonic lamina propria of infected NMRs. Without intervention, systemic inflammation associated with sepsis ensues in infected NMRs and results in high mortality. Interestingly, we demonstrate a strong therapeutic effect of probiotics comprising *Lactobacillus, Bifidobacterium, Streptococcus salvarius subsp. thermophilus* and *Enterococcus faecium* strains. Treatment with probiotics induces mucosal healing and restores intestinal homeostasis, including suppression of excessive proliferation of epithelial cells, replenishment of goblet cells and also has an anti-inflammatory effect. Taken together, we demonstrate that NMRs, beyond their use as an anti-ageing and disease-resistance model, can also be used to address disease mechanisms underlying infectious colitis, including disruptions in the mucosal barrier permeability, gut microbial ecology and in local and systemic immune regulation; and in testing functional probiotics strains as potential therapeutics.

## Introduction

Laboratory mice have been powerful tools for studying host-enteropathogen infections and underlying mechanisms of acute and chronic inflammatory diseases of the gut. These include genetically engineered mice, congenic models, chemically induced and immune cell transfer models, gnotobiotic and gut infection models amongst others^1-12^. These models have enabled the role of uncontrolled bacterial colonisation, epithelial barrier disruptions and unregulated immune stimulation to be deciphered in inflammatory diseases. Furthermore, mouse models have also been used in evaluating novel therapeutics^13-16^. There are, however, considerable differences in the innate and adaptive immunity of mice and humans, reflecting the development of divergent strategies to combat pathogenic challenges within species-specific ecological niches^17-19^. Moreover, housing laboratory mice in specific pathogen-free (SPF) conditions profoundly affects the murine basal immune state^20-24^. Similarly, there are significant differences in the presence of specific bacterial species between laboratory mice and human gut microbiota^25^. Given the taxonomic complexity, trans-kingdom interactions within the metagenome, and their role in driving local and systemic pro-inflammatory diseases, new experimental models are needed to capture the full composition and diversity of the luminal bacterial community. One of the approaches is to include other rodent species with diverse gut microbiota, immune cell repertoire and intestinal physiology closer to humans.

The naked mole rat (NMR), *Heterocephalus glaber,* is the longest-lived rodent with a maximum lifespan of >30 years^26^ that has gained much attention as an anti-cancer and anti-ageing model organism. These animals exhibit an extremely low cancer incidence^27-32^ and an attenuated decline in physiological functions with age, including in the reproductive^28^, cardiovascular^33^ and gastrointestinal systems^34^. Several aspects of NMR biology make it a highly suitable mammalian model for exploring gut homeostasis that is relevant to understanding human intestinal physiology and associated diseases. Firstly, similar to human colons^35^, NMRs also have a thicker protective mucin layer compared to mice^36,37^, suggesting that NMRs have the potential to provide a novel insight into the regulation of penetrability properties of the inner mucus and in testing novel therapeutics for enteric infections and gut inflammation in humans. We have also recently shown that the cellular kinetics of *Lgr5-*expressing crypt-based columnar cells in the NMR colon are almost identical to human *LGR5^+^* cells while being significantly slower than those observed in mice^36^. This makes NMRs ideal for studying inflammation-induced effects on intestinal stem cells and regeneration efficiency during remission in humans. NMRs also make an interesting model for studying the host-microbiota interaction, with faecal and caecal samples of these animals showing Firmicutes and Bacteroidetes as the dominant bacterial phyla which are similar to that in humans^38-40^. Even more interestingly, NMRs also possess a high load and diversity of Spirochetaceae and Mogibacteriaceae, taxa found in human populations with unique diets or exceptional longevity, such as the Hadza gatherers and supercentenarians^39,41^.

The NMR haematopoietic system has also evolved unique features, with some key differences to mice^42-45^. For example, the NMR splenic and circulating immune cell repertoire shows a higher myeloid-to-lymphoid ratio compared to mice^42^. Similar myeloid prevalence is seen in humans^17^. A higher number of granulocytes found in NMRs are mostly neutrophils, including a novel lactoferrin-high neutrophil subset that expresses several antimicrobials and is highly responsive to a bacterial-mimicking challenge^42^. Compared to their mouse counterparts, NMR macrophages have a greater phagocytic capability and produce higher pro-inflammatory cytokines^46,47^. One of the most striking aspects of the NMR immune system is the absence of natural killer cells which has been associated with the high susceptibility of this species to viral infections^42^. This likely reflects weaker viral selection pressures due to limited exposure to airborne viruses in their subterranean habitat^48,49^. Furthermore, NMRs also do not show an age-associated increase in blood leukocytes and platelets which potentially reduces chronic inflammation and delays age associated thrombosis^43^. Additionally, several inflammaging related pathways appear to be down regulated in older NMRs^43^.

Here we report the spontaneous development of bacterial-induced acute colitis in NMRs after a sporadic outbreak of *Citrobacter braakii* (*C.braakii*) infection in our captive NMR colonies. Infected NMRs displayed several clinical and histopathological features characteristic of murine^50^ and human colitis^51^, with inflammation in the mucosal and submucosal layers of the colon, accompanied by oedema, loss of goblet cells, hyperplasia and ulcerations. Treatment of NMRs with probiotics alleviated symptoms of colitis and reversed all the histopathological features in the colon, including an increase in goblet cells and reduction in hyperproliferation of crypt cells to levels seen at homeostasis in healthy NMRs. Distinct from their use in anti-ageing and anti-cancer research, our new findings highlight the potential utility of NMRs as a new experimental mammalian model in understanding epithelial barrier function defects, gut dysbiosis and inflammation triggered by enteric infections, and also in testing novel therapeutic modalities.

## Results

### Spontaneous disease outbreak in naked mole rat colonies

NMRs were housed in two non-sterile housing facilities and monitored for routine health checks in accordance with our animal lab husbandry protocols (see Methods). At one of the facilities that housed 145 naked mole rats, individuals from three separate breeding colonies (*n=9;* non-breeders) started exhibiting signs of morbidity, which included a hunched posture, reluctance to move, recumbency and skin discolouration. These affected NMRs grew progressively worse and all died within 9 weeks of when symptoms were first noticed. A week after their death, we identified another subset of NMRs (*n=7;* 5 female, 2 male) that also began displaying similar symptoms like skin discolouration and lethargy (Fig. 1A). These NMRs were immediately separated from their colonies and enrolled in a study to comprehensively characterise their disease phenotype (Supplementary Fig. 1).

**Figure 1:**
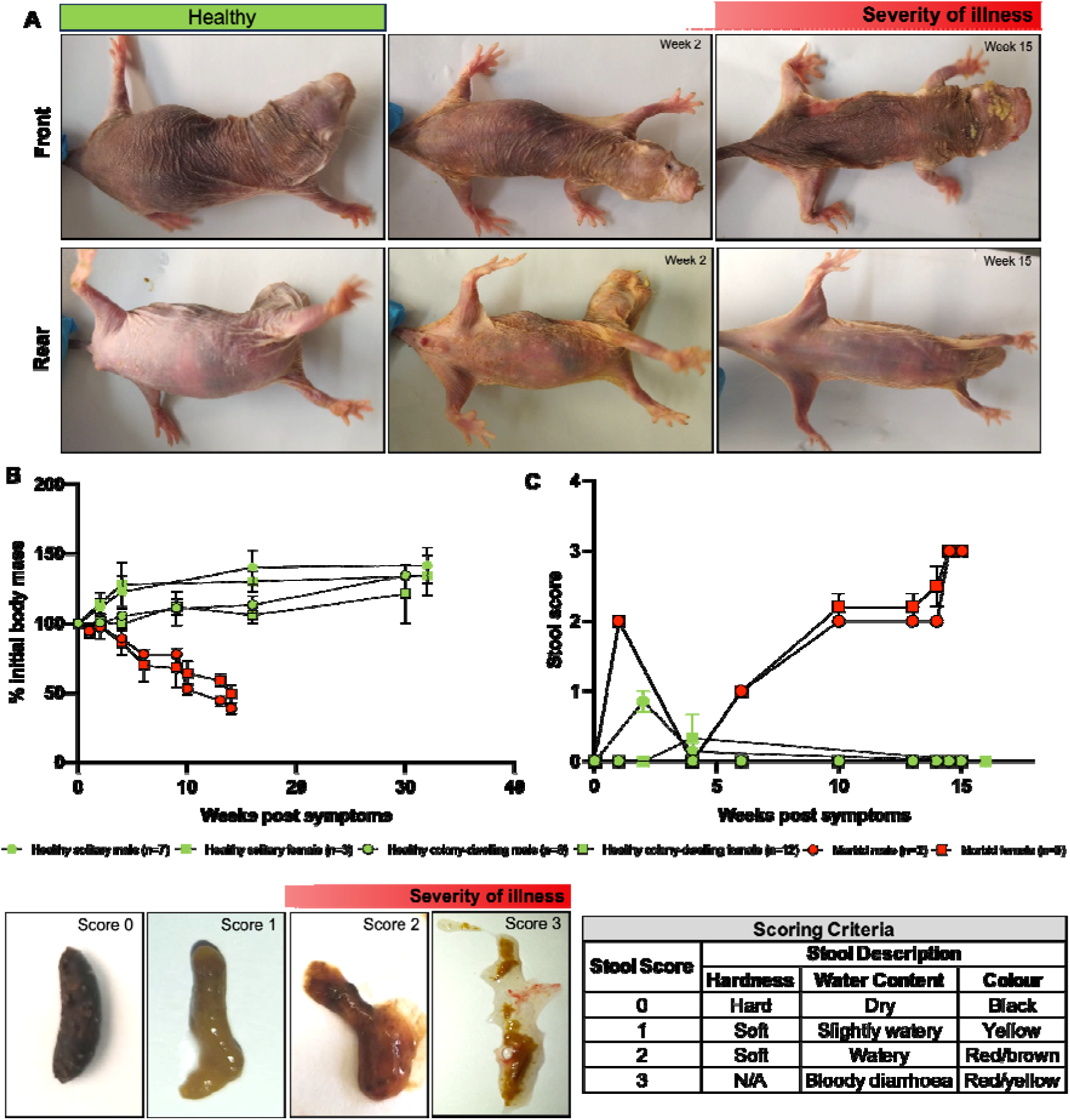
Spontaneous disease outbreak in naked mole rat colonies. (**A**) Gross morphological changes in symptomatic NMRs over 15 weeks. (**B**) Change in the body mass of morbid NMRs compared to healthy NMRs. Each data point represents the mean body mass (± SEM) measured in sick NMRs (*n*=7; 5 females (F) and 2 males (M); non-breeders; age >1year), healthy colony-dwelling NMRs (*n*=20; 12F and 8M; non-breeders; age >1year) and healthy NMRs in solitary confinement (*n=*10; 3 females and 7 males; non-breeders; age >1year). (**C**) Top, right, Graph showing changes in the stool score of sick NMRs over a 16-week period. Each data point represents the mean stool score, healthy, colony-dwelling (*n*=20), healthy, solitary (*n=*10) and sick NMRs (*n*=7). Bottom, left, Macroscopic appearance of stool samples from healthy (Score 0) to morbid NMRs with increasing disease severity (Score 1, 2 and 3). Bottom, right, Table showing the metrics used for scoring stool samples.

The separated NMRs (*n*=7) continued to deteriorate, showing additional symptoms such as rectal bleeding and severe dehydration (Fig. 1A). Rehydration with saline did not improve the condition, and by week 10, all separated NMRs lost approximately 30% of their initial body mass and then a further 25% by week 15 (Fig. 1B). In contrast, both healthy control groups, colony-dwelling (*n*=20; 12 female, 8 male) and solitary confined NMRs (*n=10*; 3 female, 7 male), continued to maintain or gain mass over the 32 weeks of observational period (Fig. 1B). We also observed a gradual change in the appearance of the stool samples in symptomatic NMRs, changing from normal, hard (Score 0) to soft, watery consistency (Score 1) by week 6 (Fig. 1C). By week 10, the stool appeared red/brown (Score 2), and by week 14, this became very watery (Score 3) (Fig. 1C). By week 14, two NMRs became immobile and presented with strained breathing. At this stage, we euthanised these animals and collected blood, liver, lung, small intestine and colon for in-depth histopathological and biochemical analysis, as discussed in the following sections. Other affected NMRs (*n*=5) displayed these severe end-stage symptoms by week 17 and were also euthanised, and tissues were collected for various assays.

### Naked mole rats are susceptible to *Citrobacter braakii* infections, a mole rat homolog of enteropathogenic *Citrobacter rodentium* in mice and *Escherichia coli* in humans

In order to characterise the cause of morbidity in NMRs, we examined the blood and various tissues from these animals. Firstly, blood smears from diseased NMRs stained with Giemsa revealed rod-shaped bacteria in both extracellular regions (Fig. 2A, left, black arrow) and within monocytes (Fig. 2A, left, blue arrow). We also found the presence of bacteria with coccoid morphology resembling *Streptococcus spp.* (Fig. 2A, right, black arrow). These observations highlighting bacteremia led us to further investigate the inflammatory responses in infected NMRs as it is well established that inflammatory signals evoked by severe systemic infections induce emergency myelopoiesis which replenishes mature blood cells^52-54^. By analysing the peripheral blood of these infected animals (*n=5*) using an automated haematological analyser^44^, we observed a 37% decrease in mature neutrophils while an increase of 5% and 28% in immature neutrophils and monocytes, respectively (Fig. 2B). In contrast, in healthy solitary NMRs (*n=4*), we observed lymphocytes and mature neutrophils as the predominant immune cell types at approximately 43 % and 52 % of the total peripheral blood cells, respectively, which is in agreement with those previously observed in NMRs housed together in colonies^44^ (Fig. 2B). There was an absence or very low presence of immature neutrophils (0%), monocytes (3%), basophils (0%) and eosinophils (0.75%) in these animals (Fig. 2B). Moreover, we observed no significant change in the proportion of lymphocytes, basophils or eosinophils in infected NMRs compared to healthy controls (Fig. 2B). Taken together, these observations show how systemic bacterial infection in NMRs causes blood-forming organs to respond by producing and migrating immature neutrophil precursors to the peripheral blood while also triggering myeloid differentiation towards monocyte lineage. Interestingly, these systemic host immune responses are in agreement with those reported in mice and human sepsis^55,56^.

**Figure 2:**
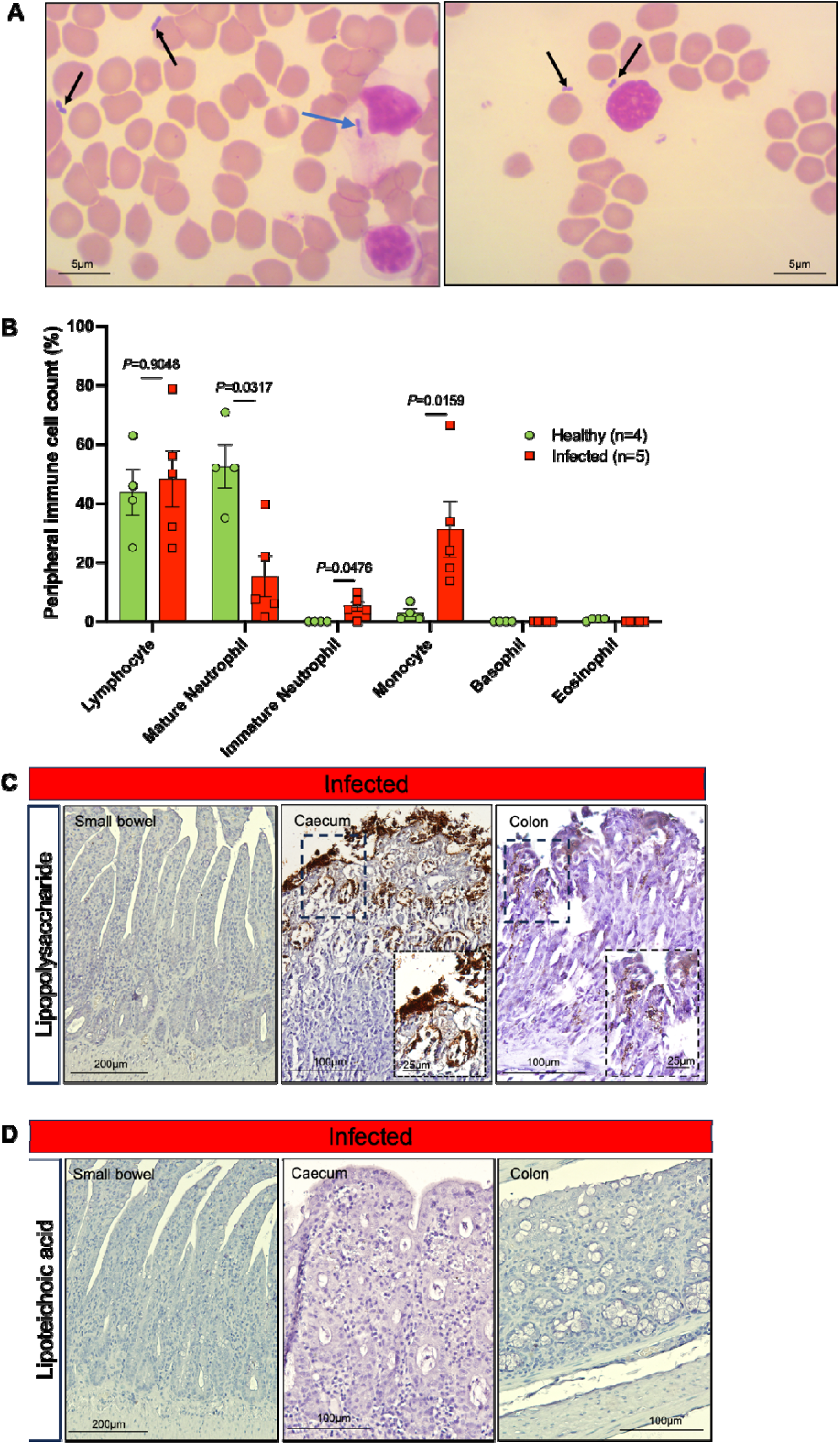
Naked mole rats (NMR) are highly susceptible to *Citrobacter braakii* infection. (**A**) Peripheral blood smear with Wright-Giemsa stain shows both (left) extracellular (black arrow) and intracellular rod-shaped bacteria (blue arrow), and (right), bacteria of coccoid morphology. (**B**) Graph showing the percentage of different immune cell populations over total leucocyte counts in the peripheral blood of healthy, solitary (*n= 4*; 1F and 3M) and infected (*n = 5*; 4F and 1M) NMRs. *P*-values generated using Mann-Whitney (two-tailed U-test) are indicated. (**C**) Photomicrographs showing small bowel, caecum and colonic tissue sections of infected NMRs stained with haemotoxylin and anti-lipopolysaccharide antibody. (**D**) Photomicrographs showing small bowel, caecum and colonic tissue sections of infected NMRs stained with haemotoxylin and anti-lipoteichoic acid antibody. Scale bars are shown as 5, 25, 100 and 200 µm.

Further histological examinations of solid tissues in infected NMR revealed multiple small foci of necrosis and inflammation in the livers (Supplementary Fig. 2A). Additionally, the lungs also displayed moderate interstitial pneumonia, characterised by mononuclear cell infiltration of the alveolar walls (Supplementary Fig. 2B). Bacterial cultures grown from affected liver samples were used in Gram staining and primary biochemical identification tests, conducted by the Laboratory Diagnostic Services at the Department of Veterinary Tropical Diseases, University of Pretoria. The analysis showed that the predominant isolate consisted of gram-negative rods, which were oxidase-negative, catalase-positive, and indole-negative, classifying them within the Enterobacteriaceae family. Finally, the API® 10 S identification system confirmed the identity of the bacterial species as *Citrobacter braakii*.

Given that the early signs of ill health (rectal bleeding, blood in stool, diarrhoea) indicated the involvement of the lower gastrointestinal tract, we assessed if the cause of bacterial septicaemia is due to the translocation of enteropathogenic bacteria from the gut into the bloodstream. To this end, we analysed the intestines of infected NMRs (*n=7*) for the presence of gram-negative and gram-positive bacteria, with anti-lipopolysaccharide (LPS) and lipoteichoic acid (LTA) antibodies. Given that we used formalin fixation and not Carnoy’s solution for tissue processing which would have preserved the mucus layer, our analysis was limited to bacterial outgrowth that had breached the mucus layer. While no LPS staining, indicative of gram-negative bacteria, was observed in the small intestine of infected NMRs, positive LPS staining was detected in close proximity to the intestinal epithelial surface of the caecum and the colon, and also in the underlying lamina propria of infected NMRs (Fig. 2C). No LTA staining was seen in the small bowel, the caecum or the colon of these infected NMRs (Fig. 2D). Taken together, these results showed gram-negative bacterial colonisation and invasion of the colon in NMRs may have led to *C. braakii* crossing the colonic mucosa and via the portal vein, producing bacterial septicaemia and subsequently liver necrosis.

To determine the relationship of *C. braakii* with other enteric bacteria, a phylogenetic tree was constructed using DNA sequences of seven housekeeping genes (adk, fumC, gyrB, icd, mdh, purA, and recA), as previously reported^57^. Using a pairwise whole-genome comparison of bacterial genomes, a high sequence homology was found between *C. braakii* and *Escherichia coli K-12* (89.54 %), *C. braakii* and *Citrobacter rodentium* (89.80 %) and *C. rodentium* and *E. coli K-12* (90.13 %). Thus *C. braakii* is closely related to members of the Enterobacteriaceae and demonstrates convergent evolution with *E. coli* (Supplementary Fig. 3). Interestingly*, C. rodentium* is a murine homolog of enteropathogenic *E. coli* and enterohaemorrhagic *E. coli* in humans and has been responsible for sporadic outbreaks of transmissible murine colitis in several laboratory mouse colonies^58-60^. Given these similarities, *C. braakii* infection could serve as a potential surrogate NMR model for enteropathogenic (EPEC) and enterohemorrhagic *E.coli* (EHEC), which poorly infect standard models like mice.

### *Citrobacter braakii i*nfected naked mole rats develop intestinal epithelial atypia

We further focused our efforts on understanding the involvement of the intestinal tissue in *C. braakii*-infected NMRs. Most of the small bowel in infected animals was largely unaffected, with mild changes detected in the most distal regions of the small intestine (Supplementary Fig. 4). In contrast, extensive architectural distortions were observed in the entire colonic mucosa of the infected NMRs (Fig. 3A). Epithelial atypia in all infected NMR colonic tissues included crypt loss in some regions (Fig. 3B) while crypt elongation in others (Fig. 3C). More specifically, crypt loss was observed in 16–60 % of the entire colon in infected NMRs and crypt elongation was detected in 11–60 % of the colon length (Fig. 3D). Other epithelial abnormalities that were observed in the majority of the infected NMRs included crypt abscesses (Fig. 3E), thickening of the muscularis propria (Fig. 3F) and crypt herniation (Fig. 3G, arrowhead). Infected NMRs also showed mucosal erosions and focal ulcerations (Fig. 3H). The extent of epithelial damage and changes in the mucosal architecture in the infected NMR colons were scored by three independent pathologists using established criteria for animal models of intestinal inflammation^6,61-63^ and is summarised in Fig. 3I. Based on the pathological scoring, surface involvement varied from 1–25 % in 14 % of infected animals to 26–50 % in 57 % of infected animals, while 29 % showed 51–75 % of surface involvement (Fig. 3I). Notably, 57 % of the infected animals showed mucosal erosions and focal ulcerations (Fig. 3I).

**Figure 3:**
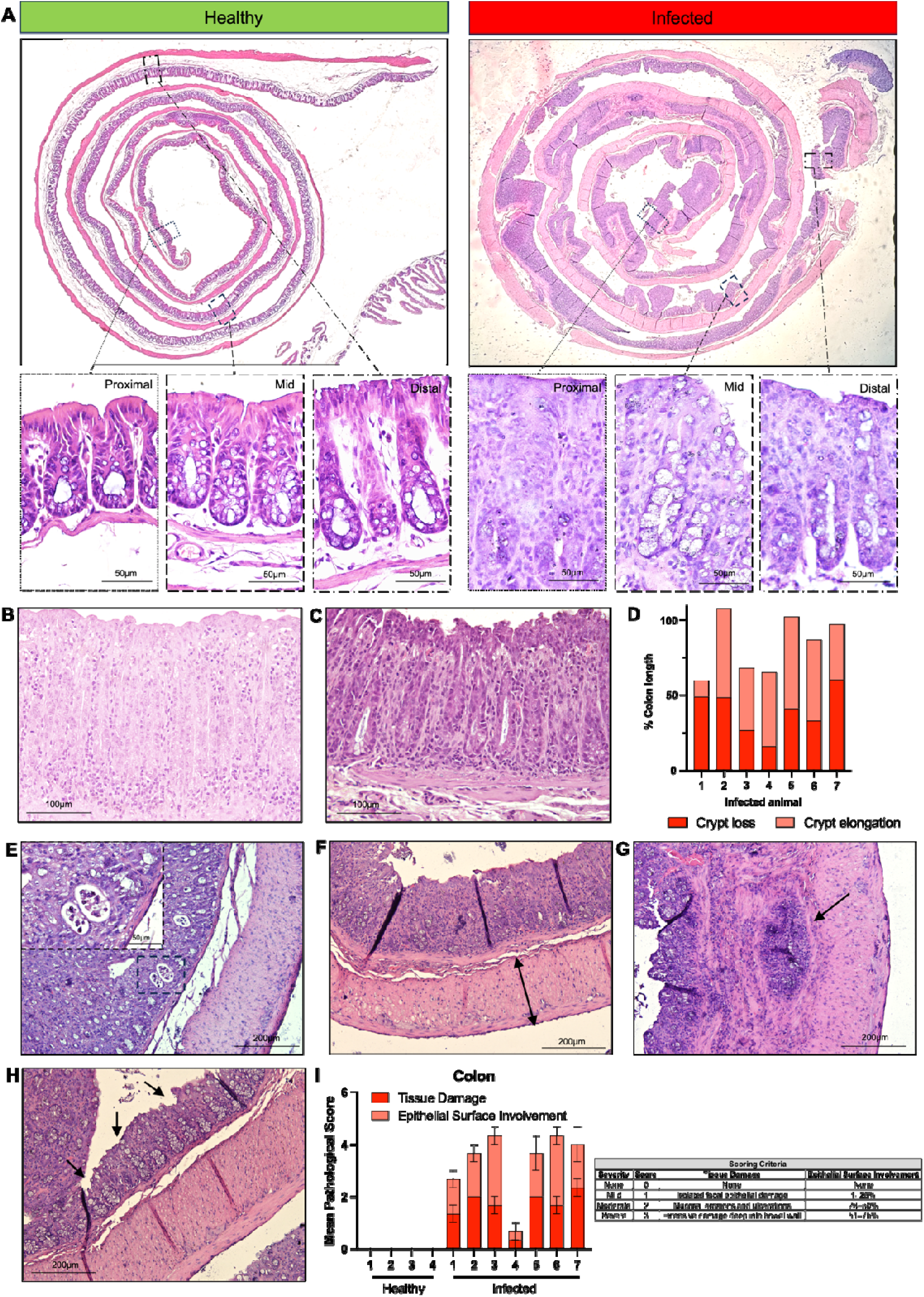
Histomorphology of colon in *Citrobacter braakii* infection in naked mole rats. (**A**) Top, Representative haemotoxylin and eosin (H&E) stained microscopic images showing colonic tissue of healthy and *C. braakii* infected NMRs. Bottom, Higher magnification images showing crypt structure in the proximal, mid and distal regions of the colon in healthy and *C.braakii* infected NMRs. **(B-H)** H&E-stained microscopic images showing aberrant epithelial histomorphology consisting of (**B**) crypt loss, (**C**) crypt elongation, (**E**) crypt abscesses, (**F**) muscularis propria thickening, (**G**) crypt herniation, and (**H**) surface erosion. (**D**) Graph showing the proportion of colonic tissue exhibiting crypt loss or elongation in *C. braakii*-infected NMRs (*n=7*). Lengths were measured on microscopic images acquired at 2X magnification using ImageJ software. (**I**) Left, Graph depicting mean pathological scores obtained from the histological assessment of infected NMRs by three independent pathologists using the scoring criteria highlighted in the table (right).

### Colonic inflammation in *Citrobacter braakii* infected naked mole rats

Due to the incompatibility of commercially available antibodies to cross-react with specific immune subtypes in the NMRs, we relied on H and E staining of infected animals to assess any tissue-specific inflammatory changes in *C.braakii*-infected NMRs. We found leukocyte infiltration in the mucosa (Fig. 4A), sub-mucosa (Fig. 4B) and also as transmural patches (Fig. 4C). Semi-quantitative evaluation method used by three independent pathologists that graded inflammation severity into mild, moderate and severe (Fig, 4D, left). This showed that 5 out of 7 infected NMRs had moderate leukocyte density within the lamina propria that was mainly localised in the mucosa and the submucosa (score 2) (Fig. 4D, right). One *C.braakii*-infected NMR showed severe inflammation which was characterised by leukocyte presence as deep as in the muscularis propria and eventually as transmural infiltrates (Score 3) (Fig. 4D).

**Figure 4:**
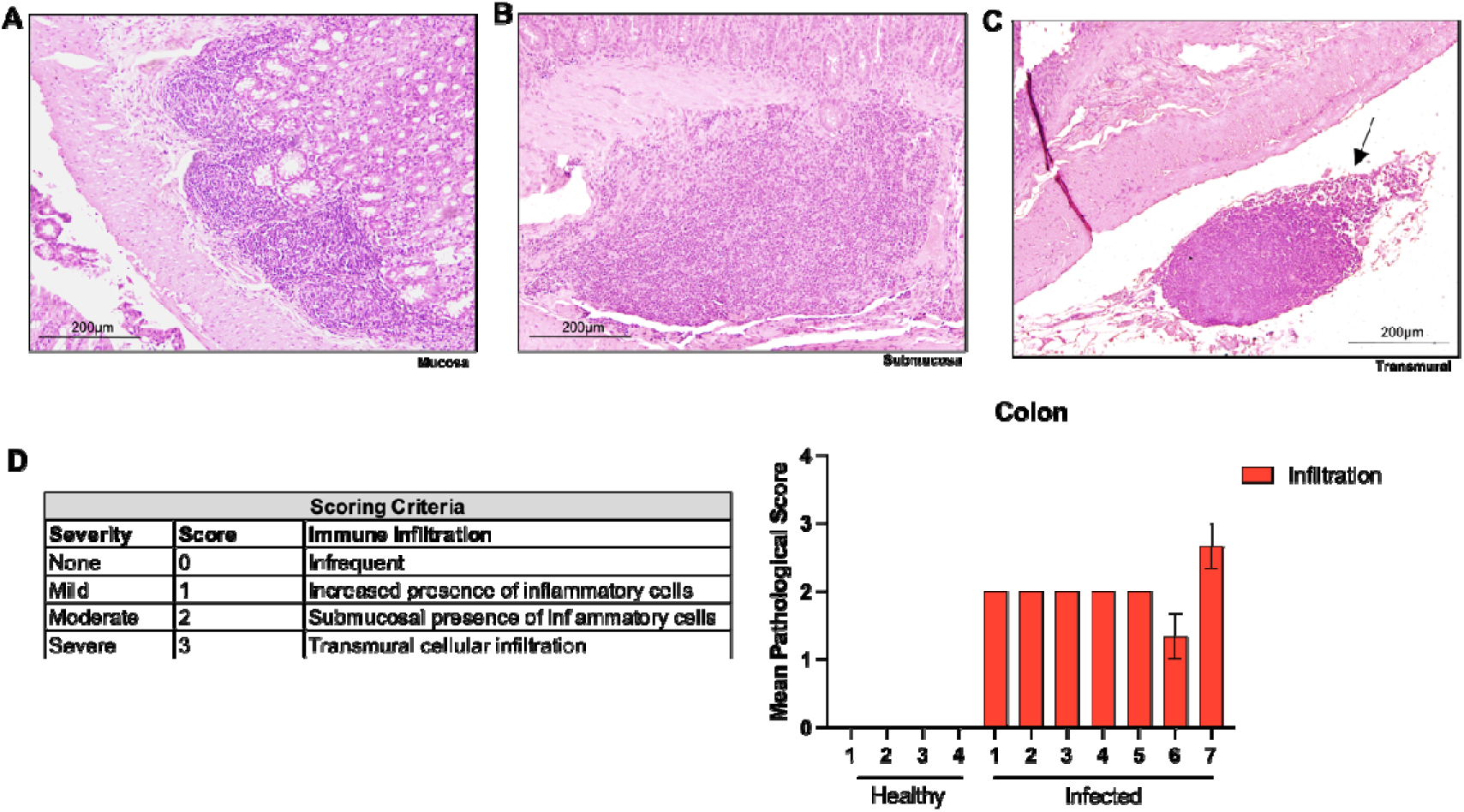
Colonic inflammation in *Citrobacter braakii*-infected naked mole rats. **(A-C)** H&E-stained microscopic image showing extensive colonic inflammation in *C. braakii* infected NMRs in (**A**) mucosal, (**B**) submucosal, and (**C**) transmural patches. (**D**) Left, Table highlighting the scoring scheme used to allow a semi-quantitative analysis of immune cell infiltrates. Right, Graph depicting the mean pathological score of inflammation in infected NMRs. Scale bars are shown as 200 µm.

### *Citrobacter braakii* infected naked mole rats show reversal of epithelial atypia after probiotics treatment

Two weeks after the second infected cohort of NMRs (*n*=7) was removed from their colony for disease characterisation, another subgroup of NMRs (*n=*10; 5 females and 5 males) also began exhibiting symptoms, including fatigue and diarrhoea (Supplementary Fig.1). We immediately isolated these animals to prevent any infectious transmission to other colony mates through, for example, coprophagy (Supplementary Fig.1). After confirmation of expansion and colonisation of *C. braakii* in the livers, colons and caecum of the first two euthanised animals from the second cohort (Supplementary Fig 1), we hypothesised that much like *C. rodentium^64^,* this pathogen might be causing infectious colitis and dysbiosis in NMRs which could be reversed by probiotic treatment. Indeed, accumulating evidence from the clinic and experimental models of inflammatory bowel disease (IBD) suggests that certain probiotic strains, especially *lactobacilli* and *bifidobacteria*, offer benefits for the treatment and prevention of IBD. We, therefore, explored the possibility of treating *C. braakii*-infected NMRs with a 7-strain probiotics cocktail (Protexin®) containing *Lactobacillus plantarum, Lactobacillus delbrueckii subsp. bulgaricus, Lactobacillus acidophilus, Lactobacillus rhamnosus, Bifidobacterium bifidum, Streptococcus salvarius subsp. thermophilus* and *Enterococcus faecium*.

We began treatment with Protexin® at different stages of the disease (mild, moderate and severe) in affected NMRs, which was determined by the time elapsed since the first observable symptoms appeared. As such, the infected were sub-divided into three treatment groups, which included NMRs displaying symptoms for 2.5 weeks (*n*=3), 6.5 weeks (*n*=3) and 11 weeks (*n*=4). Interestingly, probiotics-treated NMRs regained their strength and weight (Fig. 5A). More specifically, two groups with less established colitis (mild and moderate) gained 98% of their body mass within 3 to 4 weeks while NMRs with severe colitis took approximately 20 weeks to regain 80% of their initial body weight (Fig. 5B). Similarly, we also observed a gradual improvement in the stool score of NMRs with mild disease, changing from Score 2 to Score 0 within 3 weeks of probiotics administration (Fig. 5C). NMRs with moderate and severe colitic disease took 6 weeks of treatment to reach normal stool consistency (Fig. 5C). In addition to these external measurements of disease remission, we also euthanised NMRs from mild (*n*=1), moderate (*n*=1) and severe (*n*=1) colitis group after 48 weeks of Protexin® treatment and performed histomorphological analysis on the colons of these animals. Firstly, the length of the colon, which contracts during colitis, was similar between the treated and healthy animals (*n*=6) (Fig. 5D). The crypt structures (Fig. 5E) and muscularis propria thickness (Fig. 5F) were also similar in treated NMRs to those seen in healthy NMRs. The rest of the treated NMRs that were not euthanised continued to remain healthy (over 2 years) and have also been used in breeding protocols to start new colonies.

**Figure 5:**
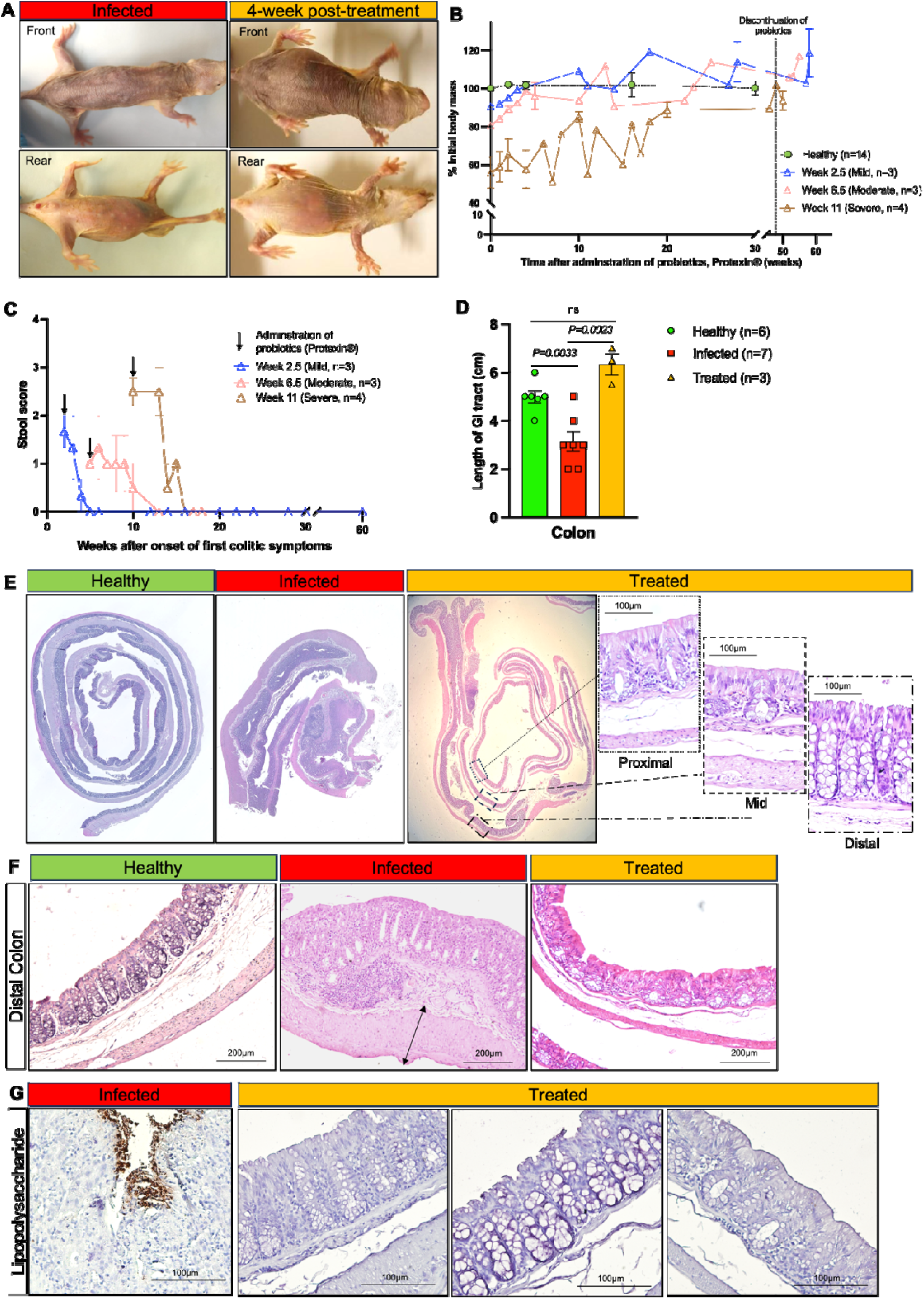
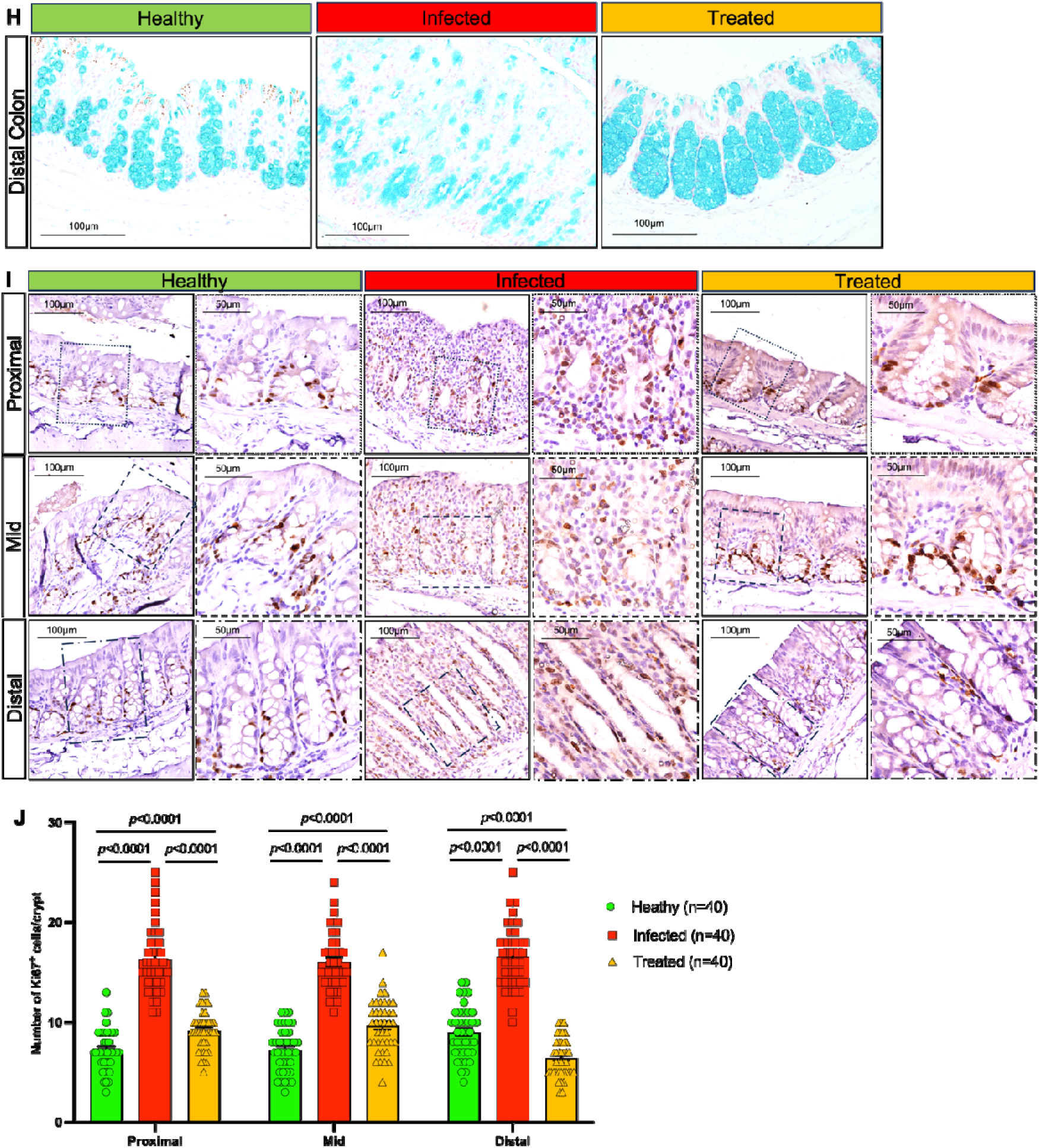
*Citrobacter braakii* infected naked mole rats show reversal of epithelial atypia after probiotics treatment. (**A**) Gross morphological changes in NMRs after 4 weeks of treatment with 7-strain probiotics (Protexin®). (**B**) Graph showing the change in the mean body mass (± SEM) of colitic NMRs treated with Protexin®. Each line represents NMRs presenting with different stages of their colitic disease. Prior to Protexin® administration, the mild group, *n*=3 (2F and 1M), had exhibited symptoms for 2.5 weeks, the moderate group of *n*=3 (1F and 2 M) for 6.5 weeks while the severe group, *n*=4 (2F and 2M), had been symptomatic for 11 weeks previously. Colony-dwelling, healthy NMRs were used as controls (*n*=14; 7F and 7M). (**C**) Change in the mean stool score (± SEM) in NMRs (*n*=10) treated at 2.5, 6.5 and 11 weeks since the onset of first symptoms. (**D**) Bar graph showing the mean length (± SEM) of colons in NMRs in healthy (*n*=*6*; 3F and 3M), infected (*n*=*7*; 5F and 2M) and in 48 weeks of probiotics-treated NMRs (*n*=3; 2F, 1M). (**E**) Representative H&E stained microscopic images showing colonic tissue of uninfected*, C. braakii* infected and probiotics-treated NMRs. Higher magnification images of the proximal, mid and distal regions of the colon in treated NMRs are also shown. (**F**) H&E stained microscopic images comparing muscularis propria thickness in uninfected, infected and probiotics-treated NMRs. Black arrow indicates muscularis propria in the infected animal. (**G**) Photomicrographs showing positive staining for lipopolysaccharide in the caecum of infected NMR compared to negative staining in the colonic tissue sections of probiotics-treated NMRs (*n*=3). (**H**) Representative Alcian Blue stained images showing goblet cells in the distal colon of healthy, *C.braakii* infected and probiotics-treated NMRs. (**I**) Immunohistochemical staining with anti-Ki67 antibody in proximal, mid and distal colons of healthy, infected and probiotics-treated NMRs. Scale bars are shown as 100 µm and 200 µm. (**J**) Bar graph showing the mean number (± SEM) of Ki67-positive cells in the proximal, mid and distal regions of the colons of healthy (*n*=2; 1F and 1M), *C. braakii*-infected (*n*=2; 2F) and probiotics-treated (*n*=2; 1F and 1M) NMRs. 40 crypts in each region of the colon were counted per animal. *P* values are indicated on the graph. In all cases, Student’s t-test using two-tailed, unpaired and unequal variance was employed. Scale bars are shown as 100 µm and 200 µm.

Positive LPS staining for gram-negative bacteria, which was evident in infected NMR colon and caecum (Fig. 2C, 5G), was no longer present in probiotics-treated NMRs (Fig. 5G). Using Alcian blue staining, we next sought to assess any changes in the abundance of goblet cells in the colon of probiotics-treated NMRs compared to healthy and infected NMRs. We found that in contrast to the lower abundance of goblet cells in infected NMRs, healthy and probiotics-treated NMRs showed a similar distribution of goblet cells per crypt (Fig. 5H). Additionally, to assess if probiotic treatment had any effect on the proliferation of cryptal cells, we performed Ki67 immunohistochemical staining on all regions of the colon in healthy, infected and treated NMRs (Fig. 5I). Quantitative analysis revealed that there was an increased number of Ki67-positive cells in the colons of infected NMRs (Fig. 5J). More specifically, compared to healthy NMRs, we found 125% more Ki67^+^ cells in the proximal colon, 112% in the mid and 83% increase in the distal colon in the infected NMRs (Fig. 5J). Notably, treatment of infected NMRs with probiotics reduced the proliferation rates of cryptal cells to homeostatic levels observed in healthy NMRs (Fig. 5J). In summary, our results showed that probiotics treatment had a profound effect in reversing the infectious colitis phenotype in NMRs and, based on these findings, all colonies were pre-emptively treated with Protexin® and no further evidence of infectious colitic disease was reported.

## Discussion

Naked mole rats have emerged as an exciting model for anti-ageing studies due to their exceptionally long lifespan, high fecundity and low burden of age-related diseases^29,65^. Much research has focussed on elucidating evolutionary pressures and biological mechanisms driving these unique traits in NMRs that ensure long-term tissue maintenance and disease resistance. However, a few reports on spontaneous and experimentally induced pathologies have highlighted specific weaknesses in the otherwise robust biology of NMRs and offer a unique opportunity to use these fascinating mammals to understand mechanisms underlying human diseases. For example, NMRs are highly susceptible to viral infections like HSV1^48^ and coronaviruses^49^, which appears to be due to relaxed viral selection pressures on NMRs as they have minimal exposure to airborne viruses while inhabiting sealed subterranean burrows. Further insight into common pathologies in NMRs has come from necropsy data in research colonies and zoo populations that have revealed the prevalence of renal tubular mineralisation, hepatic hemosiderosis, bite wounds and chronic progressive nephropathy^27,66^.

The spontaneous disease outbreak in our NMR colonies adds to the rarely reported disease processes in NMRs and highlights the critical role of intestinal homeostasis in maintaining the overall health of long-lived NMRs. Our findings serendipitously demonstrate a new role for NMRs as a highly relevant and much-needed research tool for studying the multisystem interactions of pathogenic gut bacteria in inducing local and systemic inflammation that contributes to the development of inflammatory diseases^67^ and life-threatening sepsis^68^. In the results presented here, we show that colonisation of gram-negative bacteria in the sub-mucus layers in the caecum and colons resulted in clinical and histopathological features in NMRs that closely resemble the hallmark characteristics of mouse and human colitis^50,51^. In addition to presenting with weight loss, severe dehydration, perianal ulceration and bloody diarrhoea, histomorphological analyses of the intestinal tract showed distortions in the mucosal architecture and extensive epithelial atypia. These included ulcerations, thickening of the muscularis propria, loss of surface epithelium, hyperproliferation and crypt abnormalities. We also observed goblet cell loss in the colonic epithelium, which would reduce barrier maintenance and allow increased exposure to luminal bacteria. In future studies, it will be interesting to study how goblet cell depletion is mediated by enteric infections and how this impacts colitis progression and/or host defence. Interestingly, we have previously reported that healthy NMRs have a higher abundance of goblet cells, higher gene expression levels of mucins and a thicker mucin layer in the colon compared to mice and showed that NMRs had a higher resistance to DSS-induced colitis^36^. Another recent study has shown reduced gut permeability in NMRs by measuring intestinal anion secretion induced by serotonin, bradykinin, histamine and capsaicin^37^. Despite these indications of a robust intestinal barrier in NMRs against experimentally induced irritants, we now show that natural infection by gram-negative bacteria is able to break down the epithelial integrity in the colons of these animals.

Dysregulation of immune responses in the intestinal mucosa is central to the pathogenesis of human inflammatory diseases and other forms of infectious colitis^69-71^. Indeed, bacterial-infected NMRs showed moderate to severe leukocyte infiltration specifically in the colonic mucosa and sub-mucosa. We were unable to characterise specific immune cell infiltrates in the intestinal tissue of these animals due to the incompatibility of commercially available antibodies to cross-react with NMRs. Future work using sequencing technology will need to be employed to fully characterise the immune responses associated with bacterial-induced colonic inflammation in NMRs. In addition to colonic inflammation, assessing the peripheral blood of infected NMRs highlighted a robust innate immune response systemically. More specifically, there was a significant increase in immature neutrophils and monocytes, and a concomitant decrease in mature neutrophils. These observations are in line with emergency myelopoiesis seen in septic mice^72^ or amplified granulopoiesis seen in human septic patients^55^. Interestingly, we detected both coccoid and rod-shaped bacteria in the blood and a heavy outgrowth of *Citrobacter braakii* in the livers of morbid NMRs. *C. braakii* is a gram-negative, facultative anaerobe of the *C. freundii* complex^73^ which has been found in clinical gastrointestinal specimens and is thought to be an opportunistic microorganism^74,75^. Interestingly, in mice another member of this enterobacteria family, *C. rodentium,* has been shown to induce colitis, sharing pathogenicity with human enteropathogenic *E. coli* and enterohaemorrhagic *E. coli*^64,76^. These bacteria, referred to as attaching and effacing (A/E) bacteria, share the pathogenic mechanism of colonising the intestinal mucosa by attaching to the intestinal epithelium and effacing the epithelial brush border^77-79^. Our comparative analysis revealing a high sequence homology of *C. braakii* with *C. rodentium* and *E. coli* suggests a shared enteropathogenic mechanism and related pathophysiology. Taken together, we conclude that the disease process in NMRs initiated in the lower gastrointestinal tract with *C. braakii* translocating from the caecal and colonic mucosa to the systemic circulation, resulting in liver colonisation and sepsis.

Members of the Enterobacteriaceae family, namely *Citrobacter rodentium* and *Salmonella*, have elucidated the role of enteropathogenic infections and host inflammation in disrupting colonic microbial community that provides a competitive growth advantage to the enteric pathogen and enhances its virulence^80-82^. One possible way to restore microbial balance is the use of probiotic microorganisms that have been shown to promote a healthy gut microbiota and a healthy immune system^83-86^. For example, *Bifidobacterium* spp. and *Lactobacillus* spp. probiotic treatment have been shown to reduce mortality, promote intestinal mucus secretion and reduce inflammation in in zebrafish^87^ and mice^88,89^. Similarly, VSL#3 which contains 8 bacterial strains, including *Lactobacillus, Bifidobacterium* and *Streptococcus* subspecies, has also been shown to alter intestinal permeability in *Muc2*-deficient mice and exert a beneficial effect independent of an intact mucus layer^90^. Even in humans, addition of probiotics like VSL#3 to conventional treatment of ulcerative colitis patients results in more efficacious remission^91-93^. Taking inspiration from these studies, we treated NMRs infected with *C. braakii* with a commercially available 7-strain probiotic (Protexin®). Protexin® contains *Lactobacillus plantarum, Lactobacillus delbrueckii subsp. bulgaricus, Lactobacillus acidophilus, Lactobacillus rhamnosus, Bifidobacterium bifidum, Streptococcus salvarius subsp. thermophilus* and *Enterococcus faecium*. Indeed, the use of Protexin® had a strong therapeutic effect and reversed the disease phenotype in *C.braakii*-infected NMRs. We observed restoration of the mucosal architecture, replenishment of colonic goblet cells and reduction in the proliferative index of cryptal cells that were comparable to levels seen in uninfected, healthy NMRs. While the role of individual subspecies in alleviating the effects of infectious colitis in NMRs remains to be determined, our results show how ecological rebalance in the intestinal flora by probiotics has a profound effect in promoting intestinal homeostasis in these animals.

In summary, our findings highlight how intestinal health is critical for maintaining the overall health of long-lived NMRs, and any overgrowth of a bacterial pathogen can lead to intestinal epithelial damage and pro-inflammatory responses, which culminate in bacterial-induced colitis and sepsis. Therefore, we propose using NMRs in studying specific pathogen-related colitis where new models are increasingly required to address the multifactorial aetiology and clinical heterogeneity caused by common bacteria such as *Campylobacter jejuni, Salmonella, Shigella, Escherichia coli, Yersinia enterocolitica, Clostridioides difficile,* and *Mycobacterium tuberculosis*^94^. Additionally, the strong effect of probiotics in reversing the pathophysiology of infectious colitis also underscores the use of NMRs in developing and testing microbiome-focused therapeutics against enteropathogenic infections and associated pro-inflammatory diseases of the gut.

## Methods

### Naked mole rat husbandry

Naked mole rats (NMRs) were housed in two separate, non-sterile animal facilities within the Department of Zoology and Entomology at the University of Pretoria. They inhabited non-sterile tunnel systems composed of interconnected Perspex chambers, with wood shavings provided as nesting material. All NMRs remained within their respective colonies, except when removed for breeding or experimental procedures in accordance with standard laboratory operating protocols. Animals were weighed regularly, and various biological samples, including faecal matter and blood, were collected as part of specific experimental protocols and general standard operating procedures. The ambient conditions in the NMR housing rooms were maintained at a temperature range of 29 – 32□°C, with a relative humidity of 40 – 60%, replicating the underground burrow environments of their natural habitat. NMRs were provided with a daily *ad libitum* diet consisting of chopped fresh fruits and vegetables (e.g., apple, sweet potato, cucumber, and capsicum), supplemented once daily with ProNutro (Bokomo)^95^. As NMRs obtain sufficient hydration from their food, no additional drinking water was provided. The health status of all NMRs was monitored daily.

### Isolation of symptomatic NMRs

Animals that presented with skin discolouration, weight loss, diarrhoea, bloody stool, or recumbency were removed from their colonies and housed individually in plastic chambers with sterilized wood shavings as nesting material. These animals were fed a daily *ad libitum* diet of chopped fresh fruits and vegetables (e.g., apple, sweet potato, cucumber, and capsicum) along with a bi-daily supplement of ProNutro (Bokomo).

### Probiotics treatment of symptomatic NMRs

Symptomatic NMRs that presented with skin discolouration, weight loss, diarrhoea, bloody stool, or recumbency were removed from their colonies and housed individually in plastic chambers with sterilized wood shavings as nesting material. These animals were fed a daily *ad libitum* diet of chopped fresh fruits and vegetables (e.g., apple, sweet potato, cucumber, and capsicum) along with a bi-daily supplement of ProNutro (Bokomo) mixed with 0.1 g of 7-strain probiotic cocktail (Protexin®). Protexin® contained *Lactobacillus plantarum, Lactobacillus delbrueckii subsp. bulgaricus, Lactobacillus acidophilus, Lactobacillus rhamnosus, Bifidobacterium bifidum, Streptococcus salivarius subsp. thermophilus,* and *Enterococcus faecium*.

### Tissue collection and processing

NMRs were euthanised according to approved protocols as previously described^36^. Before euthanasia, whole blood was collected. The bleed site was first cleaned by thoroughly swabbing the tail with Biotaine (0.5 % chlorhexidine gluconate, 70 % ethanol; Braun Medical, reg. no. 33/13.1/0526) and allowing it to dry. A sterile blade was then used to pierce the tail, and blood was collected using an EDTA-coated microhaematocrit capillary tube. The collected blood was immediately transferred into a sterile EDTA tube for further analysis.

Next, intestines were excised from the abdominal cavity and sub-divided into 4 segments, duodenum, jejunum, ileum and colon as reported previously^36^. All portions of the small intestine and colon were then flushed with a 1 × PBS (Phosphate Buffered Saline) solution, using a P1000 pipette to remove any faecal material. Measurements were taken to determine the length of the intestinal tract. Each segment of tissue was longitudinally incised with a gut-cutting device, and the edges were affixed to a 3 mm filter paper with the luminal side facing upwards. Tissues were fixed overnight in 10 % neutral buffered formalin at room temperature. Fixed tissues were then rolled up using the Swiss-rolling technique and transferred to 70% ethanol for storage at 4□°C. Subsequently, the formalin-fixed Swiss rolls were dehydrated through progressively increasing ethanol concentrations, cleared with xylene, and embedded in paraffin.

Liver and lung samples were excised, collected in 1XPBS and sent immediately for biochemical and pathological assessment to the Laboratory Diagnostic Services at the Department of Veterinary Tropical Diseases, University of Pretoria. Tissues were subsequently fixed in formalin and paraffin-embedded for further histological evaluation.

Finally, prior to any staining (immune or colourimetric), paraffin blocks were sectioned at a thickness of 4□µm using a microtome (Anglia Scientific) and mounted on SuperFrost Plus slides (VWR, 6310108) and dried overnight. Tissue sections were baked at 60□°C (1 h), deparaffinized by immersing slides in xylene (2 times, 10□min each) and subsequent rehydration in 100 % ethanol (2 times, 5□min each), 95 % ethanol (2□min), 70 % ethanol (2□min), 50 % ethanol (2□min), and distilled water (5□min).

### Haematoxylin and Eosin staining

Haematoxylin and Eosin staining was performed as previously described^36^. After deparaffinization and rehydration, sections were stained with Harris Haematoxylin (Merck, HHS32) (2 min 45 s) followed by a 5-minute wash under running tap water. Subsequently, slides were immersed in 95 % ethanol (10 times) before counterstaining with Eosin solution (Merck, 117081) for 3□minutes. The tissue sections were then dehydrated in 95 % ethanol (15□s) and 100 % ethanol (2 times, 15□s each), immersed in xylene (2 times, 5□min each), and finally coverslipped with DPX Mountant (Merck, 06522).

### Alcian blue staining

Alcian staining was performed as previously described^36^. Following deparaffinization and rehydration, sections were immersed in 3% acetic acid solution for 3 minutes before being stained with Alcian blue 8GX (Merck, A5268) solution (pH 2.5) for 30 minutes. The tissue sections were then washed under running tap water for 5 minutes and counterstained with Nuclear Fast Red (Merck, N3020) for 5 minutes. Following a 1-minute wash in running tap water, the tissue sections underwent dehydration in ethanol, immersion in xylene, and finally were coverslipped with DPX Mountant (Merck, 06522).

### Immunohistochemistry

Following deparaffinization and rehydration, endogenous peroxidase activity was quenched by incubating sections in 3% H_2_O_2_ (Merck, 8222871000) for 20□minutes. A heat mediated antigen retrieval was then performed by boiling sections in 1 × target retrieval solution,Citrate pH 6.0 (DAKO, S236984-2) for 10□minutes. Slides were then allowed to cool to room temperature in the same solution. Next, tissue sections underwent permeabilization through incubation with 1 × PBSTX (0.1% Triton X) for 10□minutes. All sections were then blocked for 1□hour at room temperature using 5 % serum which matched the species of the secondary antibody. Primary antibodies were prepared and diluted in antibody diluent (1 % BSA prepared in 1 × PBS) and applied to tissue sections for incubation at 4□°C, overnight in a humidified chamber. After 3 washes with 1 × PBST (0.1 % Tween20 in 1 × PBS), biotinylated secondary antibodies prepared in antibody diluent were applied to slides and incubated for 1 hour at room temperature in a humidified chamber. Target detection was performed using Avidin/Biotinylated enzyme Complex (ABC) kit (Vector Laboratories, PK-6101) and developed the signal using the DAB (3,3’-diaminobenzidine) solution (R&D systems, 4800-30-07). Lastly, tissue sections were counterstained with Harris Haematoxylin (Merck, HHS32) for 5□seconds, and underwent dehydration in 70 %, 90 % and 100 % ethanol for 15□seconds each, immersed in xylene and coverslipped with DPX Mountant (Merck, 06522). A list of all antibodies used is provided in the table below:

| Antibody | Manufacturer | Catalogue Number | Dilution |
| --- | --- | --- | --- |
| Lipopolysaccharide Core mAb WN1 | HycultBiotech | HM6011 | 1:500 |
| Lipoteichoic acid mAB | HycultBiotech | HM2048 | 1:500 |
| Ki67 | Cell Signaling | 12202 | 1:500 |

### Brightfield microscopy

Brightfield images of tissue sections were captured using an Olympus BX51 microscope coupled with an Olympus DP70 camera system using DP controller software using 4X, 10X, 20X and 40X objective lens.

### Quantification of blood immune cells

Complete blood counts were performed on NMRs whole blood samples in an automated haematological analyser (ADVIA 2120i Siemens) at the Clinical Pathology unit of the Department of Companion Animal Clinical Studies, Faculty of Veterinary Science, University of Pretoria. Briefly, blood samples were mixed with ADVIA 2120i BASO reagent which contains acid and surfactant to haemolyze the red blood cells. The white blood cells were then analysed using two-angle laser light scatter signals to obtain the total leucocyte counts. Next, the analyser used the Peroxidase method to classify and quantitatively measure leucocyte cell count and leucocyte cell type, namely, neutrophils, lymphocytes, monocytes, eosinophils, large unstained cells, and basophils, based on cell-specific constituents when the cells are treated with peroxidase stain.

### Wright–Giemsa staining

Blood smears from EDTA blood were prepared on glass slides and subsequently stained with a Wright-Giemsa stain using an automated stainer (Aerospray Hematology Pro, EliTech Group, USA). Photomicrographs were captured using a Zeiss Axio Lab A1 microscope with a Zeiss Axiocam ERc5s, using Zeiss Zen software.

### Ph2ylogenetic analysis

The sequences for each of seven housekeeping genes (adk, fumC, gyrB, icd, mdh, purA, and recA) were extracted from the following enteric bacteria: *Citrobacter braakii* strain MiY-A (accession number: NZ_CP045771), *Citrobacter rodentium* ICC168 (accession number: NC_013716.1), *Citrobacter koseri* ATCC BAA-895 (accession number: NC_009792.1), *Cronobacter sakazakii* ATCC BAA-894 (accession number: NC_009778.1), *Escherichia coli* O127:H6 str. E2348/69 (accession number: NC_011601.1), *Escherichia coli* O157:H7 str. Sakai (accession number: NC_002695.2), *Escherichia coli* str. K-12 substr. MG1655 (accession number NC_000913.3), *Klebsiella variicola* (strain: LEMB11) (accession number: NZ_CP045783.1), *Pectobacterium atrosepticum* SCRI1043 (accession number: NC_004547.2), *Salmonella enterica* subsp. *enterica serovar* Enteritidis str. P125109 (accession number: NC_011294.1), *Salmonella enterica* subsp. *enterica* serovar Typhi str. CT18 (NC_003198.1), *Salmonella enterica* subsp. *enterica* serovar *Typhimurium* str. LT2 (accession number: NC_003197.2), Yersinia enterocolitica subsp. enterocolitica 8081 (accession number: NC_008800.1), and individually aligned using MUSCLE^96^ and subsequently concatenated. MEGA11 were used to identify the best phylogenetic model to fit the data using maximum likelihood algorithm. Phylogenetic tree was constructed by the maximum-likelihood method using the recommended General Time Reversible (GTR) plus Gamma model distributed with invariant sites. The reliability of the tree was estimated with the bootstrap method with replicates set at 500 and was rooted using an outgroup comprising Yersinia and Pectobacterium. Genome sequence homology was performed using Basic Local Alignment Search Tool (BLAST).

### Statistical Analysis

All quantitative data are presented as mean□±□SEM. Statistical differences between two experimental groups were analyzed by two-tailed unpaired Student’s t-test or Mann–Whitney test using GraphPad Prism 10.2.0. The levels of significance for individual data sets are indicated in the corresponding figure legends.

## Acknowledgements

Special thanks to Dr Yolandi Rautenbach, Carien Muller and the rest of the Clinical Pathology team at the Department of Companion Animal Clinical Studies, Faculty of Veterinary Science, University of Pretoria for conducting the haematological analysis for this project. We also extend our gratitude to Mr. Erick Kapp from the Bacteriology Laboratory, Department of Veterinary Tropical Diseases, Faculty of Veterinary Science, University of Pretoria, for performing the bacteriology tests. We are also thankful to Prof Nan Gao (Rutgers, The State University of New Jersey), Dr Sandrine Ménard (Institut de Recherche en Santé Digestive, Université de Toulouse) and Sir Walter Bodmer (University of Oxford) for their comments on the manuscript.

## Author Contributions

Naked mole rats and ethics for in vivo work were acquired by N.C.B. and D.W.H. Animal experiments were conducted by D.W.H. Histological staining, microscopy, phylogenetic and statistical analyses were performed by A.S.N. Histopathological scoring was performed by P.G., R.G. and P.G. Post-mortem examination was done by N.O. Tissue processing was done by S.M. Intellectual input was provided by D.W.H., N.K., J.E.E. and I.P.M., and N.C.B. Funding was acquired by J.E.E, N.C.B and S.I. S.I. wrote the manuscript with feedback from all of the other authors.

## Animal Ethics

All scientific procedures involving NMRs were carried out with approval from the Animal Ethics Committee at the University of Pretoria under the licenses EC034-18, NAS046-19, NAS199-2020, NAS289-2020, NAS017-2021 and NAS022-2021. Additionally, approval from the Department of Agriculture, Forestry, and Fisheries (DAFF) under section 20 was obtained (SDAH-Epi-20111909592, SDAH-Epi-21021810221 and SDAH-Epi-21021810222, SDAH-Epi-20072707041).

## Patient and public involvement

Patients and/or the public were not involved in the design, or conduct, or reporting, or dissemination plans of our research.

## Supplementary Figures

**Supplementary Figure 1:**
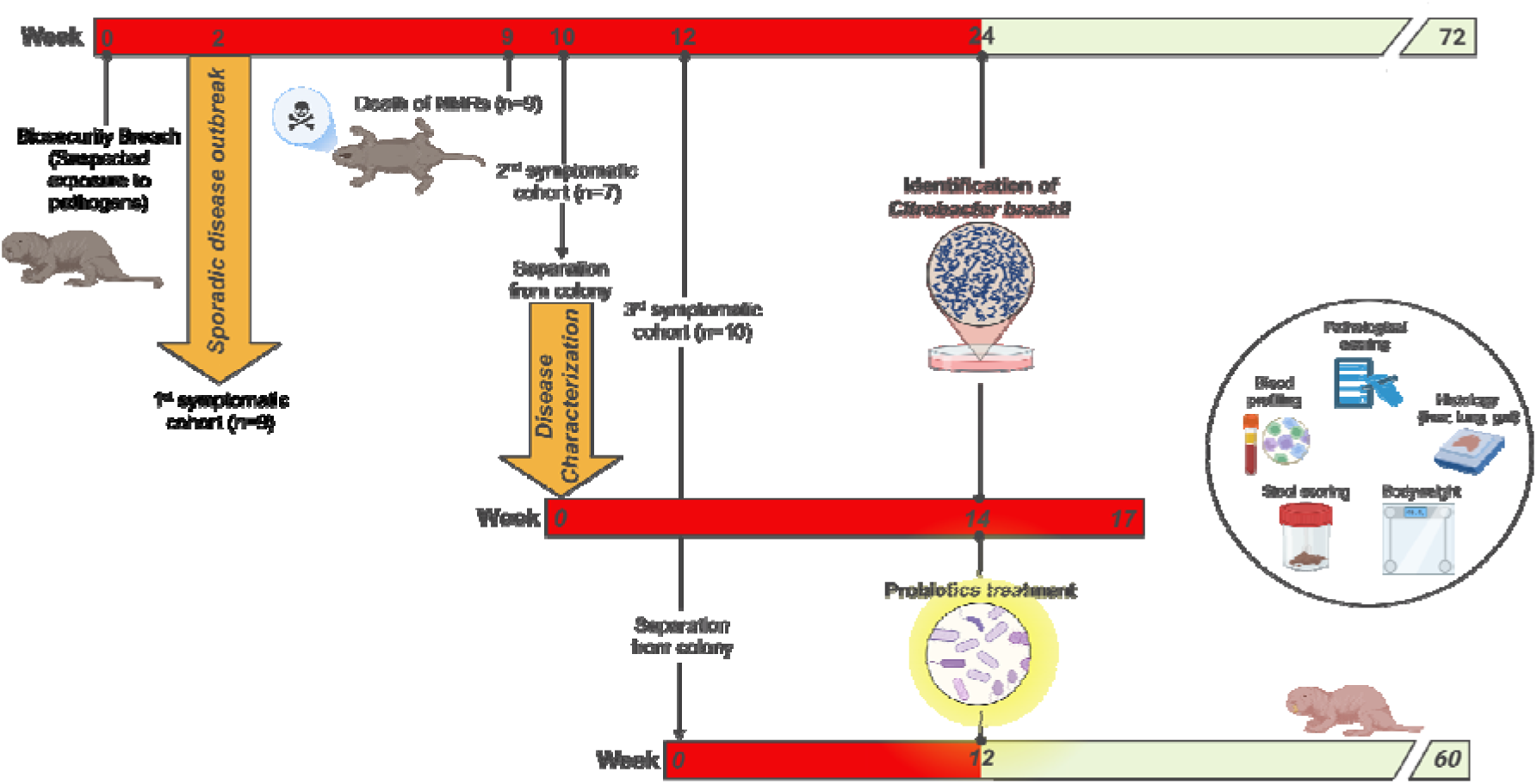
A schematic representation of the sporadic disease outbreak in our naked mole rat colonies that were observed over a period of 72 weeks after a suspected poor biosecurity practice resulted in pathogen transmission and disease onset. The first symptoms were detected in a subset of NMRs (*n*=9) which subsequently succumbed to disease-induced mortality. A week later, a second cohort of NMRs (*n*=7; 5F and 2M) showed similar symptoms and were immediately separated from their colonies. These animals were used for disease characterisation over a 17-week period, which included blood sampling, stool sample collection, body mass measurements and histopathological scoring of liver, lung and intestinal samples. Biochemical testing of diseased tissue led to the identification of *Citrobacter braakii* as the pathogen responsible for the outbreak. A third diseased cohort (*n*=10; 5F and 5M) were treated with 7-strain probiotics (Protexin®) for 48 weeks, and stool samples, body mass, blood immune profiling and intestinal histological analysis performed on these animals.

**Supplementary Figure 2:**
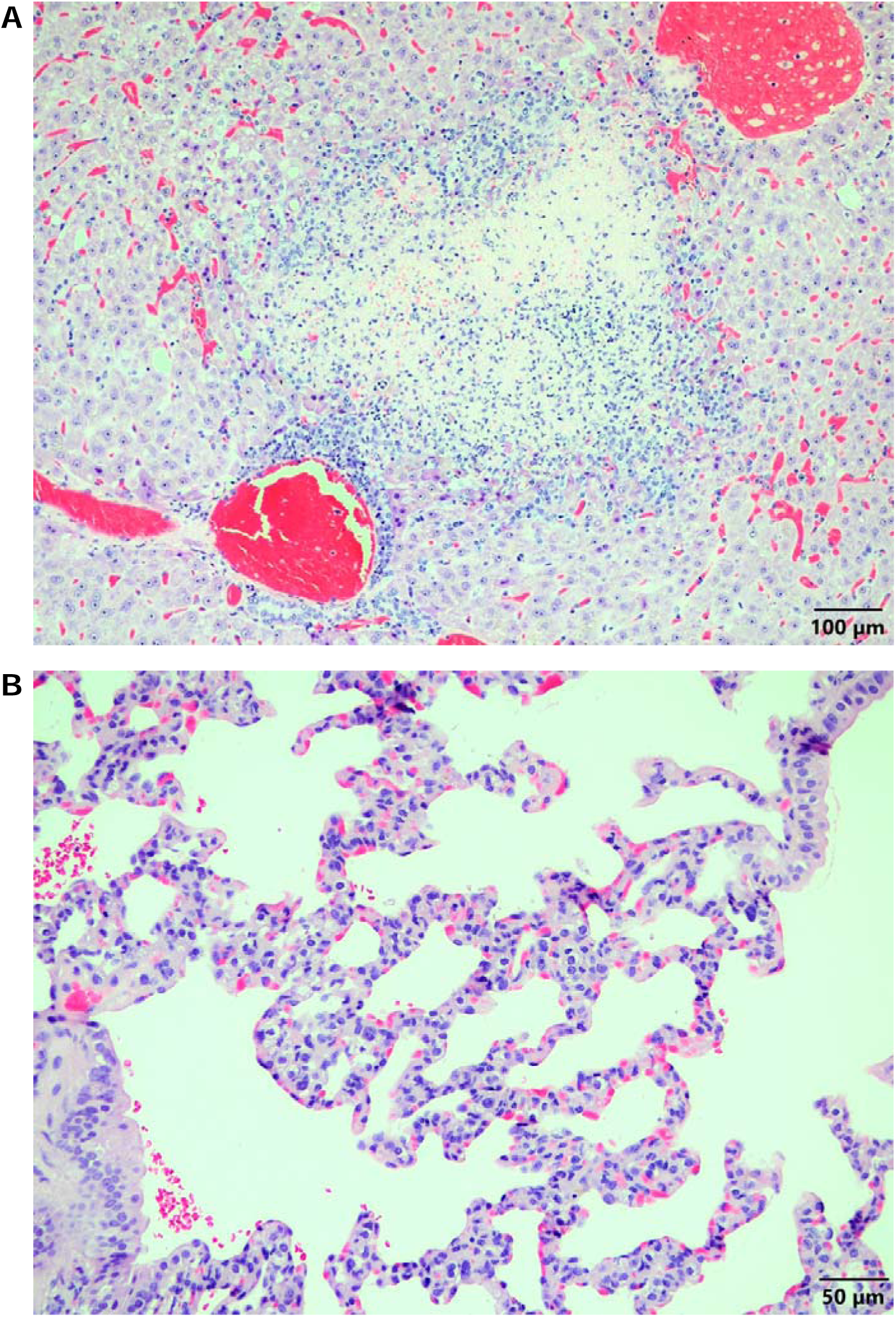
(A) H&E-stained microscopic image of liver tissue from symptomatic NMR showing multifocal foci of necrosis of varying sizes, randomly distributed across the tissue. Many of these foci were located near or superimposed with portal triads. The centres of the necrotic foci exhibited lytic necrosis, surrounded by lymphocytes, macrophages, and occasional neutrophils. **(B)** H&E-stained microscopic image of lung tissue from another diseased NMR showing multifocal foci of necrosis and moderate interstitial pneumonia associated with mononuclear cell infiltration of the alveolar walls. Scale bars are shown as 50 µm and 100 µm.

**Supplementary Figure 3:**
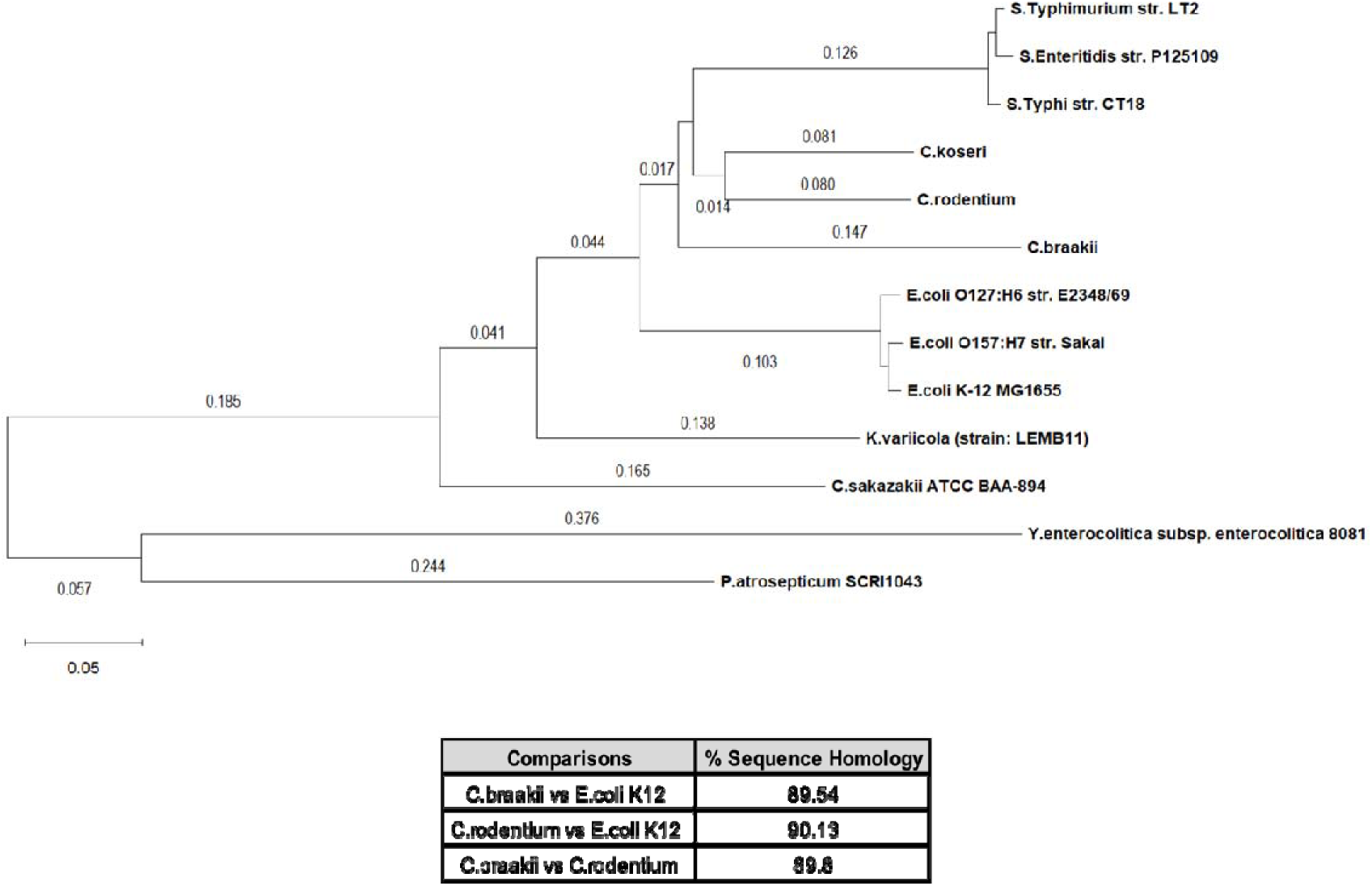
Top, Phylogenetic relationship of *C. braakii* to various enteric bacteria based on the nucleotide sequences of seven housekeeping genes (adk, fumC, gyrB, icd, mdh, purA, and recA). Bottom, Genome sequence homology of *C. braakii* to various enteric bacteria.

**Supplementary Figure 4:**
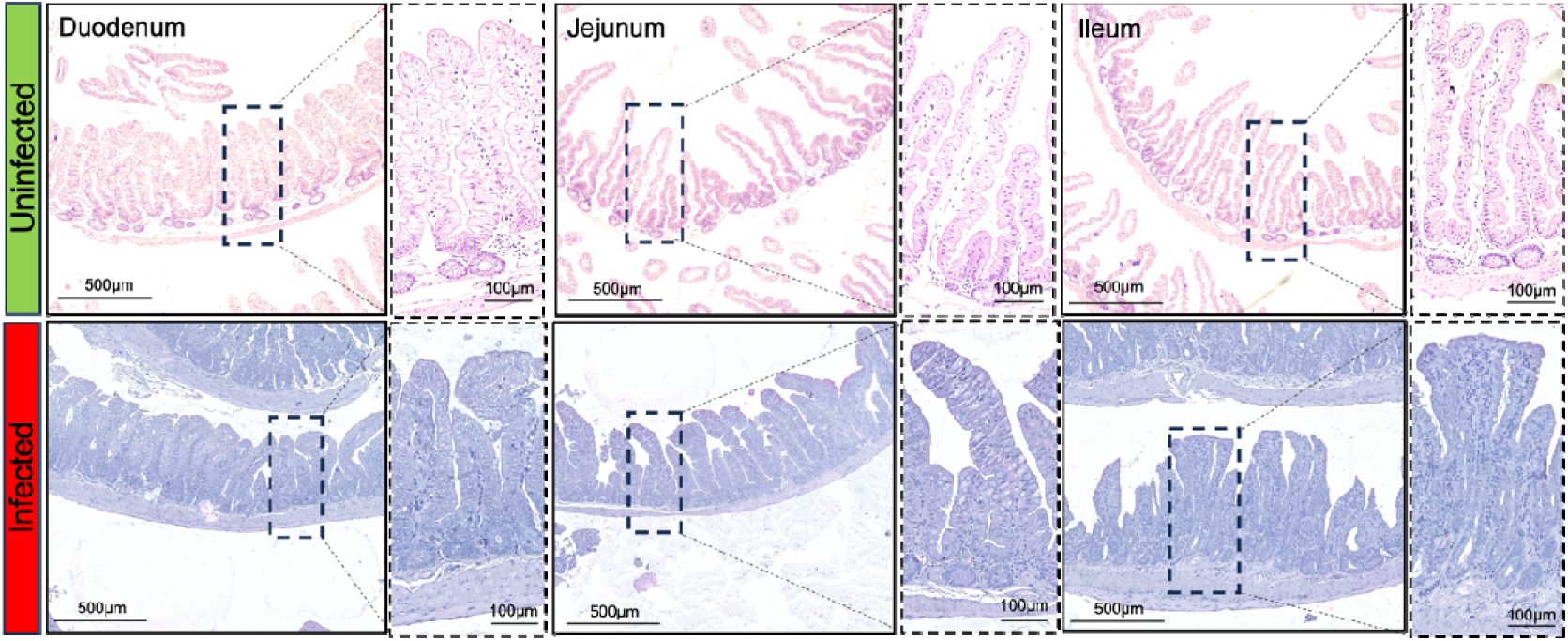
Representative H&E stained microscopic images showing small bowel tissue of uninfected and *C. braakii* infected NMRs with higher magnification images in the proximal, mid and distal regions of the colon. Scale bars are shown as 100 µm and 500 µm.

